# Stress-induced vascular remodeling: novel insight into the role of omega-3 fatty acid metabolite, 4-oxoDHA

**DOI:** 10.1101/2023.07.25.550603

**Authors:** Makoto Nishimori, Naomi Hayasaka, Kazunori Otsui, Nobutaka Inoue, Manabu Nagao, Ryuji Toh, Tatsuro Ishida, Ken-ichi Hirata, Tomoyuki Furuyashiki, Masakazu Shinohara

## Abstract

**Background:** Stress has garnered significant attention as a prominent risk factor for inflammation-related diseases, particularly cardiovascular diseases (CVDs). However, the precise mechanisms underlying stress-driven CVDs remain elusive, thereby impeding the development of effective preventive and therapeutic strategies.

**Methods:** To explore the correlation between plasma lipid metabolites and depressive states, we conducted a study involving healthy volunteers (n=408). Liquid chromatography (LC)/mass spectrometry (MS)/MS-based lipidomics and the self-rating depression (SDS) scale questionnaire were employed for data collection. In addition, we utilized a mouse model by subjecting mice to restraint stress and investigating the impact of stress on plasma lipid metabolites and vascular remodeling following carotid ligation. In vitro functional and mechanistic studies were performed using macrophages, endothelial cells, and neutrophil cells.

**Results:** Our findings revealed a significant association between depressive state and reduced plasma levels of 4-oxoDHA, a specific omega-3 fatty acid metabolite regulated by 5-lipoxygenase (LO) in neutrophils in healthy volunteers. In mice, restraint stress led to decreased plasma 4-oxoDHA levels and exacerbated vascular remodeling. Moreover, 4-oxoDHA demonstrated the ability to enhance Nrf2-HO-1 pathways, exerting anti-inflammatory effects on endothelial cells and macrophages. Mechanistically, stress-induced noradrenaline triggered the degradation of 5-LO in neutrophils through the proteasome system, facilitated by dopamine D2-like receptor activation. The reduction in circulating 4-oxoDHA resulted in the downregulation of the Nrf2-HO-1 anti-inflammatory axis and an increase in ICAM-1 expression, vascular permeability, and remodeling.

**Conclusions:** Our study unveiled a novel stress-induced pathway of vascular inflammation, mediated through the regulation of omega-3 fatty acid metabolites. Reduced levels of circulating 4-oxoDHA under stress conditions may serve as a promising biomarker for stress. This understanding of the interplay between neurobiology and lipid metabolism provides a potential avenue for the development of treatments aimed at preventing stress-induced systemic neuroinflammation.

**Highlights:**

– Our study reveals that stress-induced reduction in circulating levels of a specific omega-3 fatty acid metabolite, 4-oxoDHA, contributes to vascular inflammation.
– We have identified a novel pathway that explains how stress promotes systemic vascular inflammation by regulating omega-3 fatty acid metabolites in the circulation.
– Our findings provide new evidence for the role of 4-oxoDHA in maintaining Nrf2-ARE-related anti-inflammatory functions in endothelial cells and macrophages.

## Introduction

Stress refers to the physiological and psychological responses that occur in the presence of prolonged or excessive stimulation from various internal and external stressors. While stress responses serve as adaptive mechanisms for maintaining homeostasis, chronic or severe stress can lead to mental and physical health problems such as depression, anxiety, and cognitive dysfunction, thereby increasing the risk of illness (1, 2). Notably, the physiological response to stress influences immune, endocrine, and metabolic pathways, with stress-induced imbalances often leading to inflammation. Recently, stress has received increased attention as a substantial and modifiable risk factor for various inflammation-related diseases, including cardiovascular diseases (3–7).

Uncontrolled inflammation is recognized as a common underlying factor in many chronic diseases. Exploring intrinsic regulatory mechanisms within the inflammatory response can offer new insights into disease pathogenesis and potential treatment approaches. The acute inflammatory response is protective and involves various lipid mediators, including eicosanoids (prostaglandins and leukotrienes) derived from the essential fatty acid arachidonic acid, as well as cytokines and chemokines (8–12). Lipids play critical roles in cellular functions, including the regulation of protein transport, anchoring, and structural support. In neuronal function, lipids are essential for processes such as membrane fluidity, permeability, vesicular formation and transport, neurotransmitter release, cell integrity, and plasticity (13, 14). The levels of low-density lipoproteins, absolute polyunsaturated fatty acids (PUFAs), and PUFA subtype ratios have been proposed as potential biomarkers for major depressive disorder (15). Omega-3 PUFAs have demonstrated efficacy in improving depression (16). Considering stress as a significant risk factor for depression and stress-related physical disorders like cardiovascular diseases, investigating how omega-3 PUFAs alleviate psychosomatic symptoms becomes crucial. Therefore, our focus in this study is on PUFA-derived metabolites in the plasma. While several studies have identified potential blood biomarkers for stress, such as proteins, monoamine metabolites, and lipids, these findings have often involved cross-sectional comparisons between control and diagnosed major depressive disorder groups (*15*). Thus, there is a need to establish sensitive early diagnostic biomarkers for the pre-symptomatic stage of stress. In this study, we investigated the correlation between depressive states and plasma lipid metabolites in healthy volunteers. Our findings revealed a correlation between depressive states and specific omega-3 fatty acid metabolites in the plasma. Moreover, we validated these observations in a mouse model of stress, which resulted in enhanced vascular remodeling. To the best of our knowledge, these findings uncover a novel pathway through which stress promotes systemic vascular inflammation by regulating omega-3 fatty acid metabolites in the circulation.

## Methods

### Reagents

RAW 264.7, HL-60, EA.hy 926, and HUVEC were obtained from the American Type Culture Collection (ATCC, Manassas, VA, USA), Riken BRC (BioResource Research Center, JP), and Lonza (Basel, Switzerland). RAW264.7 and EA.hy 926 cells were cultured in Dulbecco’s Modified Eagle’s medium (DMEM). HL-60 cells were cultured in Roswell Park Memorial Institute (RPMI)-1640 with 1 µM of all-trans retinoic acid from Wako (Wako Pure Chemical Industries Ltd., Osaka, JP). HUVEC cells were cultured in Endothelial Cell Growth Medium (Lonza). All other media were purchased from Wako and supplemented with 10% fetal bovine serum (FBS; Gibco by Life Technologies, Grand Island, NY, USA). For specific agonism and antagonism of each catecholamine receptor, we employed following reagents: DL-norepinephrine hydrochloride (Sigma Aldrich), cirazoline hydrochloride (α1 adrenergic agonist, TOCRIS), B-HT 933 dihydrochloride (α2 adrenergic agonist, TOCRIS), DL-isoproterenol hydrochloride (non-selective β adrenergic agonist, Sigma Aldrich), dobutamine hydrochloride(β1 adrenergic agonist, TCI), clenbuterol hydrochloride (β2 adrenergic agonist, Sigma Aldrich), BRL 37344 sodium salt hydrate (β3 adrenergic agonist, Sigma Aldrich), phentolamine mesylate (non-selective α adrenergic antagonist, Selleck Chemicals), propranolol hydrochloride (non-selective β adrenergic antagonist, Selleck Chemicals), dopamine hydrochloride (Wako), SKF81297 hydrobromide (D1-like receptor agonist, TOCRIS), quinpirole hydrochloride (D2-like receptor agonist, TOCRIS), SCH 39166 (D1-like receptor antagonist, TOCRIS), raclopride (D2-like receptor antagonist, Funakoshi), and barbadin (β-arrestin inhibitor, Cosmo Bio Co., Ltd.).

### Clinical participants and health checkups

The clinical study was approved by the ethical review board of Kobe University Medical Ethical Committee (No. 781) on September 19, 2017. All clinical participants were healthy volunteers who had undergone regular health check-ups. Residual plasma samples collected during health checkups were frozen and stored at -80°C. The self-rating depression scale (SDS), designed by Zung (17), was used to score the depressive state of participants, which was then classified into four categories of depression severity: within the normal range (below 40 points), minimal to mild depression (40–47 points), moderate to marked depression (48–55 points), and severe to extreme depression (56 points and above).

### Restraint stress mouse model

All animal experiments were approved by the Animal Care and Use Committee of Kobe University (permission number: P190802). Six-week-old adult male C57BL/6J mice (weighing 20–25 g) were housed in standard cages at a temperature of 20–25°C under a 12-h day/night cycle in pathogen-free conditions with ad libitum access to food and water. The mice were randomly assigned to stress treatment groups. Stressed mice were restrained for 2 h in a confined space that prevented them from moving freely or turning around but did not unduly compress them. This method induces chronic stress, as evidenced by neuroendocrine activation and induction of anxiety and depression, but does not cause pain or injury (18, 19).

### Vascular remodeling in mouse carotid artery

We used a mouse model of vascular remodeling by carotid ligation (20). After general anesthesia by intraperitoneal injection of a combination of midazolam, butorphanol, and medetomidine, the right carotid arteries were dissected and entirely ligated at the carotid bifurcation. Mice were euthanized seven days after ligation and then perfusion-fixed under physiological pressure with 10% neutral-buffered formalin.

### Morphological analysis

The vessel was removed and embedded in a Tissue-Tek optimal cutting temperature (OCT) compound (Sakura, Japan), and serial sections (5 μm) were cut for histological analysis using hematoxylin and eosin (HE). For immunohistochemistry, sections were stained with anti-CD45 antibody (1:100, BD Biosciences, Franklin Lakes, NJ, USA #550539) and anti-ICAM-1 antibody (1:100, Abcam, #ab119871), followed by incubation with Takara POD conjugate solution and DAB substrate (Takara, Japan).

### LC/MS/MS-based lipidomics

To quantify lipid mediator levels in plasma, 1 mL of iced methanol per 100 µL of plasma was added and incubated for 1 h to extract total lipids. To quantify lipid mediator levels in bone marrow, whole bone marrow was flushed directly with 100% methanol at -20 °C (21). All samples for LC-MS-based lipidomics were extracted using solid-phase extraction columns. Deuterated internal standards, d4-LTB_4_, d8-5-HETE, d4-PGE_2_, and d5-RvD_2_, representing each chromatographic region of the identified LMs, were added to the samples (500 pg each) to facilitate quantification. The samples were extracted by SPE on C18 columns, as previously described (22), and subjected to LC-MS/MS. The system consisted of a Q-Trap 6500 system (Sciex) equipped with a Shimadzu LC-30AD HPLC system. A ZORBAX Eclipse Plus C18 column (100 mm × 4.6 mm, 3.5 µm, Agilent Technologies) was used. The mobile phase used was methanol/water/acetic acid at a gradient of 55:45:0.01 to 98:2:0.01 (v/v/v) at a 0.4 ml/min flow rate. To monitor and quantify the levels of the targeted LMs, the multiple reaction monitoring (MRM) method was developed with signature ion pairs Q1 (parent ion)/Q3 (characteristic fragment ion) for each molecule. Identification was conducted according to published criteria using the LC retention time, specific fragmentation patterns, and at least six diagnostic fragmentation ions. Quantification was performed based on the peak area of the MRM chromatograph, and linear calibration curves were obtained with authentic standards for each compound.

### RNA sequencing

Total RNA was extracted from HUVEC using the RNeasy Mini Kit Plus (QIAGEN) and quantified using a NanoDrop (Thermo Fisher Scientific). RNA-seq was performed after a quality check (RIN value >9.0) by Rhelixa Co. (Tokyo, JP) using an Illumina NovaSeq 6000 (Illumina, Inc., CA, USA). For pre-processing, the Trimmomatic toolkit (23) was used to trim the sequences and remove short low-quality reads, and Salmon (24) was used to quantify transcript abundance from RNA-seq reads. Gene clustering (t-SNE) and pathway analyses were performed using the iDEP toolkit (Integrated Differential Expression and Pathway Analysis ver. 0.96) (25).

### Protein analysis by western blotting

Total cell protein lysates were prepared by homogenizing the cells with radioimmunoprecipitation assay (RIPA) buffer, followed by sonication. Total protein concentration was measured using the Bicinchoninic Acid (BCA) protein assay. An aliquot of total cell lysate corresponding to 10 µg was electrophoresed using 10% sodium dodecyl sulfate-polyacrylamide gel electrophoresis (SDS-PAGE), transferred onto a polyvinylidene difluoride (PVDF) membrane, and blotted using an iBlot 2 Dry Blotting System (Invitrogen, Massachusetts, USA). Blotting membranes were blocked with 5% skim milk in Tris-buffered saline with Tween-20 for 30 min and subsequently incubated with a primary antibody diluted in Signal Enhancer HIKARI Solution (Nacalai Tesque). Antibodies against rabbit anti-5-lipoxygenase (1:1000, Cell Signaling, #C49G1), rabbit anti-NRF2 (1:500, Cell Signaling #D129C), rabbit anti-heme oxygenase-1 (1:2000, Abcam, #ab52957), mouse anti-beta-actin (1:10000, Abcam, #ab3013), and rabbit anti-PCNA (1:1000, Cell Signaling, #D3H8P) were used. After washing, the membranes were incubated with horseradish peroxidase-conjugated horse anti-mouse IgG (1:10,000; Cell Signaling, #7076S) or goat anti-rabbit IgG antibody (1:10,000; Cell Signaling, #7074S). Immunoreactivity was detected using an Immobilon Western chemiluminescent horseradish peroxidase substrate (Millipore Corporation, Billerica, MA, USA) and visualized using a ChemiDoc Touch Imaging System (Bio-Rad Laboratories Inc., Hercules, CA, USA). The volumetric value of each band was assessed using ImageJ software. To investigate the amount of Nrf2 transcription factor in the nuclear fractions, a HUVEC culture corresponding to one million cells was collected. Nuclear fractions were obtained using NE-PER Nuclear and Cytoplasmic Extraction Reagents (Thermo Scientific, IL, USA).

### Nrf2 ARE-binding activity assay

The ARE-binding activity of Nrf2 was measured in nuclear extracts using a commercially available kit (Nrf2 Transcription Factor Assay Kit, No. 600590, Cayman Chemical Company, Michigan, USA) following the manufacturer’s instructions.

### Endothelial cell permeability assay

EA.hy 926 cells were seeded into a transwell chamber with a 0.4 µm pore polyester membrane insert (Corning) at a density of 1 × 10^5^ cells/chamber two days before the treatment. After incubation with the indicated lipids for 1 h, H_2_O_2_ (500 µM)-induced permeability was quantified using FITC-conjugated dextran (10 kDa, 1 mg/mL, Tokyo Chemical Industry Co., Ltd., JP) for 2 h. The fluorescence intensity in the lower chambers was determined using a fluorescence reader (EnSpire Multimode Plate Reader, PerkinElmer, Waltham, MA, USA).

### Flow cytometry

Cell analyses were performed using an Attune Acoustic Focusing Cytometer (Thermo Fisher Scientific). For the cell viability assay, HL-60 cells were collected after incubation with noradrenaline (10–1000 µM) for 2.5 h and stained with Annexin V-FITC and PI (Annexin V-FITC Apoptosis Detection Kit, Nacalai Tesque, JP) for 15 min at room temperature.

For quantification of 5-lipoxygenase protein expression in mouse neutrophils, mouse whole blood (100 µL) was incubated with 1 × RBC lysis buffer (Invitrogen, eBioscience) on ice for 5 min, fixed and permeabilized with Cytofix/Cytoperm solution (BD Biosciences, CA, USA) on ice for 20 min, and incubated with 5-LO monoclonal antibody (Abcam, 1:100, #ab169755) on ice for 30 min. Cells were then washed twice with Cytoperm/Cytowash solution, followed by incubation with anti-rabbit IgG and Alexa Fluor 488-labeled secondary antibodies (1:100, Bioss Antibodies, Massachusetts, USA) on ice for 30 min. The neutrophil fraction was determined using FFC/SSC scatter plotting, and the expression levels of 5-LO were measured.

### RT-PCR analysis

Total RNA was extracted using an RNeasy Mini Kit (Qiagen, Hilden, Germany). cDNA was prepared from 1 µg of total RNA using the PrimeScript RT Reagent Kit with gDNA Eraser (Takara). Real-time polymerase chain reaction (real-time PCR) was performed with TB Green Premix Ex Taq II (Takara). Primers were obtained from Takara Bio Inc. Amplification reactions were performed in duplicate using a LightCycler 96 Real-Time PCR system (Roche Molecular Systems, CA, USA), and fluorescence curves were analyzed using this software. 18S ribosomal RNA was used as an internal control. Relative quantification was performed using the ΔΔCt method.

### Macrophage phagocytosis

RAW 264.7 cells (0.5 × 10^5^) were adhered onto 96-well plates and incubated with or without 1 µM DHA, 4-HDHA, or 4-oxoDHA for 6 h at 37 °C. Cells were then incubated with fluorescent-labeled opsonized zymosan (Molecular Probes) for 30 min at 37 °C. The plates were gently washed, extracellular fluorescence was quenched by the addition of 1:40 diluted trypan blue solution, and phagocytosis was determined by measuring the total fluorescence intensity using a fluorescent plate reader (EnSpire, PerkinElmer).

### Statistical analysis

Results are expressed as the mean ± standard error (s.e.). Statistical significance was determined using a two-tailed Student’s t-test for two-group comparisons and one-way analysis of variance (ANOVA) for multiple comparisons, with post hoc analysis using Tukey’s test (GraphPad Prism). Statistical significance was set at p<0.05.

## Results

### Depressive state correlates with low plasma 4-oxoDHA levels in healthy volunteers

We first investigated whether a depressive state correlated with plasma fatty acid metabolite profiles in healthy volunteers. The depressive state was quantitatively assessed using the SDS score obtained by answering twenty questions (17). Plasma was processed for liquid chromatography (LC)/mass spectrometry (MS)/MS-based lipidomics analysis. A total of 408 samples were analyzed, of which 54.1% were obtained from males. The mean [standard deviation] age and SDS score were 54.1 [11.7] and 37.2 [8.0] years. All monohydroxy fatty acids are expressed as the percent conversion ratio of each substrate. Among the variables, including lipid profiles, age, and sex, 4-oxoDHA (coefficient, - 60.941; p = 0.002), age (coefficient, -0.1; p = 0.003), and 18-HEPE (coefficient, -83.671; p = 0.024) significantly correlated with SDS scores in univariate analysis (Table 1, left). In multiple regression analysis, age (coefficient, -0.093; p = 0.006) and 4-oxoDHA (coefficient, -52.013, p = 0.019) significantly correlated with SDS scores. These epidemiological data indicated that a depressive state correlated with low plasma 4-oxoDHA levels in healthy volunteers.

**Table 1.**
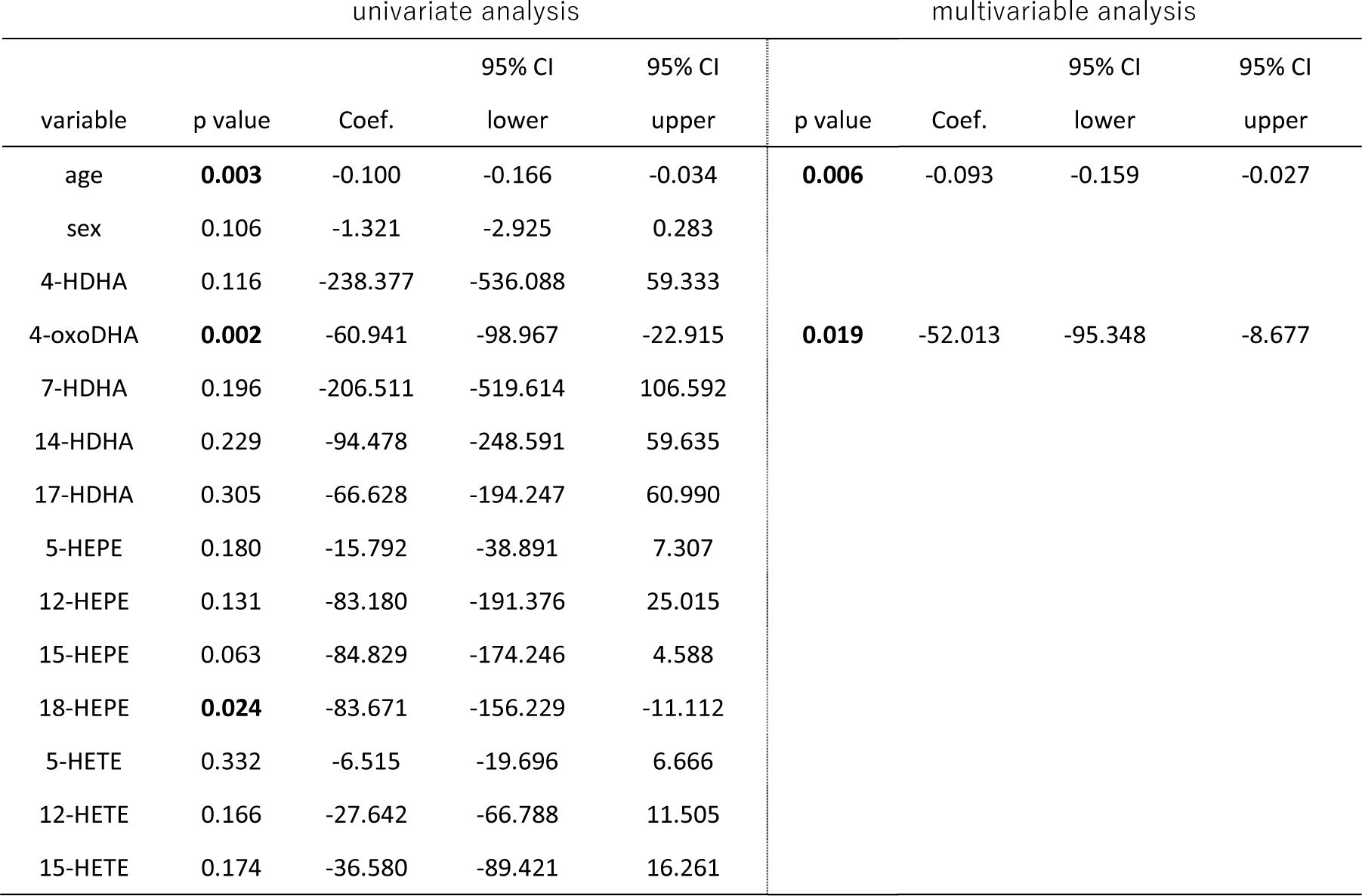
A depressive state correlates with low plasma 4-oxoDHA levels in healthy volunteers. All monohydroxy fatty acids are expressed as the percent conversion ratio of each substrate. Among the fatty acid metabolites, 4-oxoDHA independently correlated with SDS after adjusting for age, sex, and other fatty acid metabolites (n=408).

| univariate analysis |  |  |  |  | multivariable analysis |  |  |  |
| --- | --- | --- | --- | --- | --- | --- | --- | --- |
| variable | p value | Coef. | 95% CI<br>lower | 95% CI<br>upper | p value | Coef. | 95% CI<br>lower | 95% CI<br>upper |
| age | **0.003** | -0.100 | -0.166 | -0.034 | **0.006** | -0.093 | -0.159 | -0.027 |
| sex | 0.106 | -1.321 | -2.925 | 0.283 |  |  |  |  |
| 4-HDHA | 0.116 | -238.377 | -536.088 | 59.333 |  |  |  |  |
| 4-oxoDHA | **0.002** | -60.941 | -98.967 | -22.915 | **0.019** | -52.013 | -95.348 | -8.677 |
| 7-HDHA | 0.196 | -206.511 | -519.614 | 106.592 |  |  |  |  |
| 14-HDHA | 0.229 | -94.478 | -248.591 | 59.635 |  |  |  |  |
| 17-HDHA | 0.305 | -66.628 | -194.247 | 60.990 |  |  |  |  |
| 5-HEPE | 0.180 | -15.792 | -38.891 | 7.307 |  |  |  |  |
| 12-HEPE | 0.131 | -83.180 | -191.376 | 25.015 |  |  |  |  |
| 15-HEPE | 0.063 | -84.829 | -174.246 | 4.588 |  |  |  |  |
| 18-HEPE | **0.024** | -83.671 | -156.229 | -11.112 |  |  |  |  |
| 5-HETE | 0.332 | -6.515 | -19.696 | 6.666 |  |  |  |  |
| 12-HETE | 0.166 | -27.642 | -66.788 | 11.505 |  |  |  |  |
| 15-HETE | 0.174 | -36.580 | -89.421 | 16.261 |  |  |  |  |

### Restraint stress reduces plasma 4-oxoDHA levels and deteriorates vascular remodeling in mice

We next investigated stress-induced alterations in plasma 4-oxoDHA levels in mice. After 2 h of restraint stress in mice for two consecutive days, we profiled DHA metabolites in the plasma on day 3 (Figure 1a). The concentration of 4-oxoDHA significantly decreased in plasma after two days of restraint, a trend similar to that of 4-oxoDHA levels in human plasma in a depressive state. We hypothesized that the DHA metabolite 4-oxoDHA, which is decreased by stress, might have specific bioactive functions.

**Figure 1.**
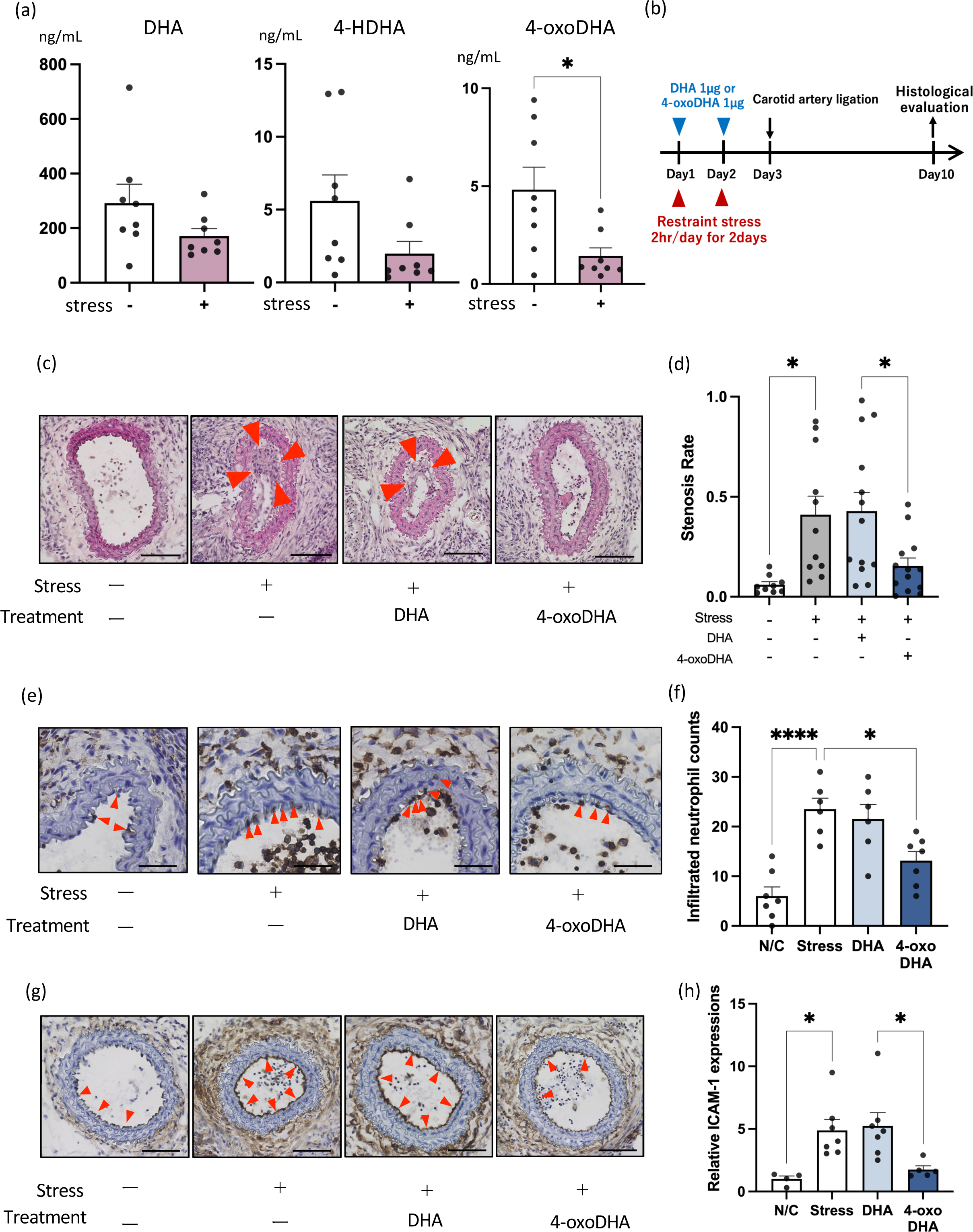
Restraint stress reduces plasma 4-oxoDHA levels and deteriorates vascular remodeling in mice. (a) Mice were treated with or without stress by physical restriction (2h) for two consecutive days. DHA, 4-HDHA, and 4-oxoDHA levels in plasma were analyzed using LC/MS/MS on day 3. Data are shown as ng/mL in plasma, mean±s.e. (n=8). * indicates p<0.05. (b) Experimental protocol for carotid artery ligation model. Mice were treated with or without stress as above; then right common carotid artery was subjected to the carotid ligation model (see method) on day 3. Specific mice were treated with 1 µg of DHA or 4-oxoDHA (intraperitoneal injection) on days of restraint stress. On day 10, vascular remodeling was histologically evaluated. (c) Representative sections at 300 µm from the carotid bifurcation are shown. Scales indicate 100 µm. (d) Stenosis rate (neointimal lesion area/lumen area). Data are expressed as mean±s.e. (n=9–12). * indicates p<0.05. (e) Infiltration of neutrophils (arrowheads) across endothelial cells was evaluated by immunohistochemistry with an anti-CD45 antibody. Scales indicate 50 µm. (f) Number of infiltrated neutrophils per section. *, **** indicate p<0.05, p<0.0001. (g) ICAM-1 expression in endothelial cells at 500 µm from the bifurcation was evaluated by immunohistochemistry with an anti-ICAM-1 antibody. Scales indicate 100 µm. (h) Relative ICAM-1 expression was quantified using Image-J software. Data are expressed as fold change compared to control, mean±s.e. (n=4–7). * indicates p<0.05.

To evaluate how stress regulates vascular remodeling, we employed a mouse carotid ligation model. As shown in Figure 1b, mice were treated with or without restraint stress; then, the right common carotid artery was subjected to carotid ligation on day 3. On day 10, vascular remodeling was histologically evaluated. We found that stressed mice showed a ∼7-fold increase in neointimal formation (arrowheads, Figure 1c), quantified by stenosis rate (Figure 1d). Next, we investigated the effects of transperitoneal administration of DHA or 4-oxoDHA on neointimal formation in stressed mice. We injected DHA (1 µg) and 4-oxoDHA (1 µg) on days of restraint stress (Figure 1b) and found that 4-oxoDHA treatment rescued the deterioration of vascular remodeling (Figure 1c, 1d).

We investigated the infiltration of neutrophils across endothelial cells by immunohistochemistry using an anti-CD45 antibody (Figure 1e, arrowheads). The number of infiltrated neutrophils was quantified (Figure 1f). We found that stressed mice showed an approximately 5-fold increase in neutrophil infiltration, and treatment with 4-oxoDHA significantly decreased neutrophil infiltration under stressed conditions.

Adhesion molecules are crucial for initiating vascular remodeling in carotid ligation models. Therefore, we immunohistochemically evaluated the induction of adhesion molecules in endothelial cells (Figure 1g, arrowheads). We found that ICAM-1 expression increased ∼5-fold in stressed mice (Figure 1h) and that 4-oxoDHA treatment almost entirely rescued stress-induced augmentation of ICAM-1.

Taken together, restraint stress reduced plasma 4-oxoDHA levels and deteriorated vascular remodeling via augmentation of ICAM-1 expression and neutrophil infiltration, and treatment with 4-oxoDHA rescued these deteriorations.

### 4-oxoDHA augments Nrf2-HO-1 pathways and anti-inflammatory properties in endothelial cells

Because 4-oxoDHA can reduce the expression of adhesion molecules in vivo, we investigated the biological functions of 4-oxoDHA in endothelial cells in vitro. First, to comprehensively examine the effects of 4-oxoDHA on HUVEC, HUVEC were incubated with or without 10 µM 4-oxoDHA for 2 h, followed by RNA-sequencing. Pathway analysis revealed that the Nrf2-ARE regulatory pathway was activated after treatment with 4-oxoDHA (Figure 2a). Next, we examined the protein expression levels of heme oxygenase-1 (HO-1), as HO-1 is significantly regulated by Nrf2-ARE. HUVEC were treated with DHA (10 µM), 4-HDHA (0.1-10 µM), or 4-oxoDHA (0.1-10 µM) for 6 h at 37 °C. Whole-cell lysates were processed for western blot analysis to determine HO-1 expression. We found that HO-1 expression levels showed a ∼7-fold increase following 4-HDHA treatment (10 µM) and a ∼28-fold increase following 4-oxoDHA treatment (10 µM) (Figure 2b).

**Figure 2.**
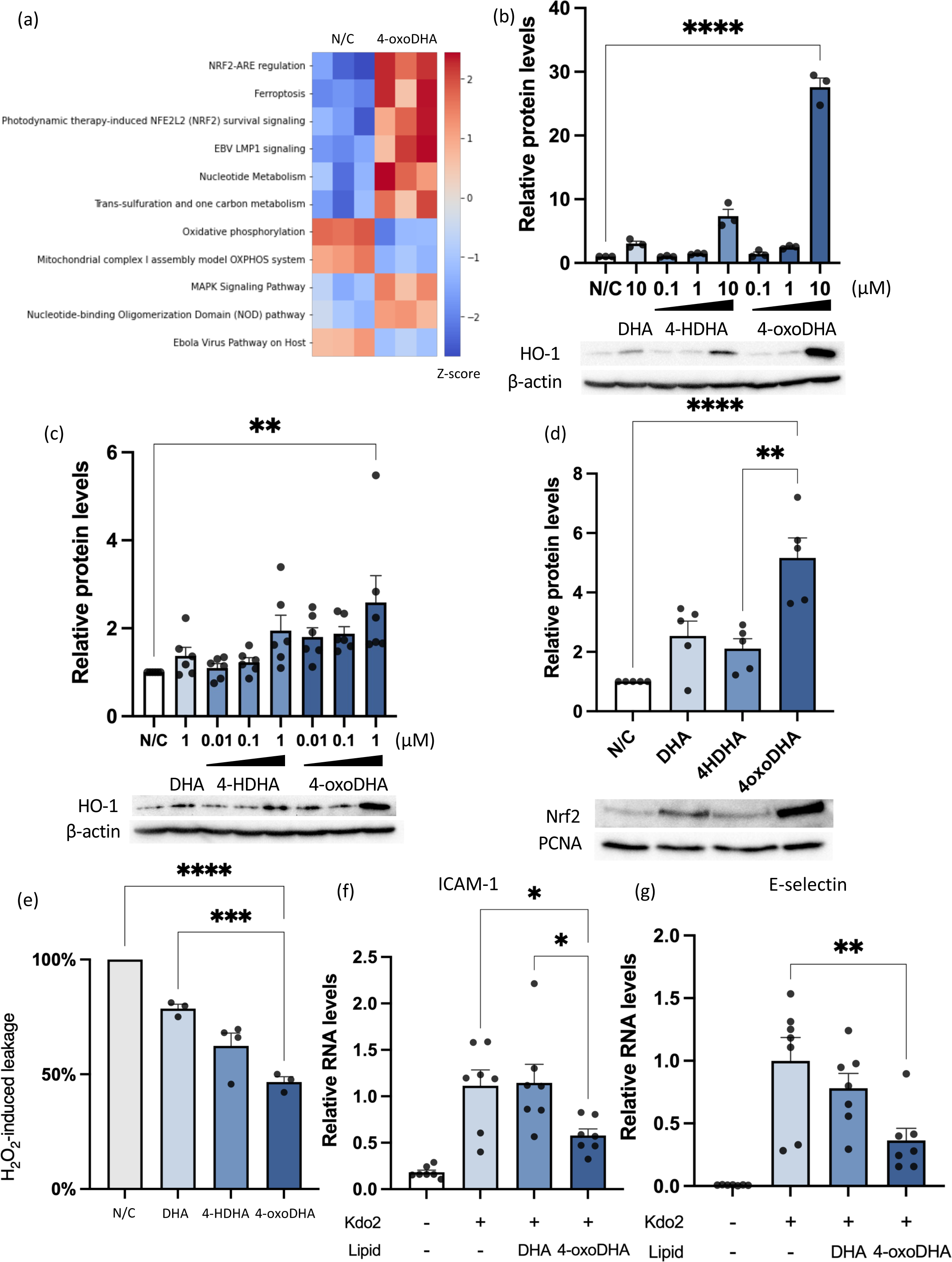
4-oxoDHA augments Nrf2-HO-1 pathways and anti-inflammatory properties in endothelial cells. (a) Pathway analysis of RNA-sequence data from the control (N/C) and 4-oxoDHA treated (4-oxoDHA) HUVECs. (b) HUVECs were treated with DHA (10 µM), 4-HDHA (0.1–10 µM), and 4-oxoDHA (0.1–10 µM) for 6h at 37°C. Whole-cell lysates were processed for western blot analysis to determine heme oxygenase-1 (HO-1) expression. β-actin was employed as the internal standard. Data are expressed as fold induction of HO-1 compared to control, mean±s.e. (n=3). **** indicates p<0.0001. (c) Endothelial cell line (EA.hy 926) was treated with DHA (1 µM), 4-HDHA (0.01–1 µM), and 4-oxoDHA (0.01–1 µM) for 6h at 37°C. Whole-cell lysates were processed for western blot analysis to determine HO-1 expression. β-actin was used as the internal standard. Data are expressed as fold induction of HO-1 compared to control, mean±s.e. (n=5). ** indicates p<0.01. (d) HUVECs were treated with 10 µM of DHA, 4-HDHA, and 4-oxoDHA for 2h at 37°C. Nuclear fractions were processed for western blot analysis to determine Nrf2 expression. PCNA was used as the internal standard. Data are expressed as fold induction of Nrf2 compared to control (N/C), mean±s.e. (n=3) **, *** indicate p<0.01, p<0.001. (e) EA.hy 926 cells were treated with 1 µM of DHA, 4-HDHA, and 4-oxoDHA for 1h, followed by stimulation of H_2_O_2_ (500 µM) for 2h at 37°C. After incubation, endothelial permeability was investigated with FITC-labeled dextran (MW: 10,000). Data are expressed as percent reduction compared to control, mean±s.e. (n=3–4). ** indicates p<0.01. (f, g) HUVECs were treated with 10 µM DHA or 4-oxoDHA after stimulation with Kdo2 (0.5 µg/mL). ICAM-1 (f) and E-selectin (g) mRNA were analyzed using real-time RT-PCR. Data are expressed as fold change compared to control, mean±s.e. (n=7). *, ** indicate p<0.05, p<0.01.

We also confirmed 4-oxoDHA-induced HO-1 expression in an endothelial-like cell line, EA.hy 926, and found that HO-1 was augmented by 4-oxoDHA treatment (Figure 2c). Next, we investigated the activation of Nrf2 in HUVEC and found that nuclear Nrf2 protein levels showed a ∼5-fold increase with 4-oxoDHA treatment (Figure 2d).

As barrier function plays a critical role in endothelial cells, we investigated the effect of 4-oxoDHA on the vascular permeability of endothelial cells. EA.hy 926 cells were treated with 1 µM DHA, 4-HDHA, or 4-oxoDHA for 1 h, followed by stimulation with H_2_O_2_ (500 µM) for 2 h at 37 °C. After incubation, we investigated endothelial permeability with FITC-labeled dextran and found that 4-oxoDHA rescued approximately 50% of vascular leakage induced by H_2_O_2_ (Figure 2e).

4-oxoDHA regulated endothelial adhesion molecules in vivo (Figure 1g, 1h). We investigated the expression of adhesion molecules in HUVEC using real-time RT-PCR. We treated HUVEC with 10 µM of DHA or 4-oxoDHA after stimulation with Kdo2-Lipid A, a TLR4 ligand (0.5 µg/mL), and found that Kdo2-induced ICAM-1 (Figure 2f) and E-selectin (Figure 2g) mRNA were reduced to 1/2 and 1/3 after 4-oxoDHA treatment.

These results indicate that 4-oxoDHA augments the Nrf2-HO1 pathway and has anti-inflammatory properties in endothelial cells.

### 4-oxoDHA augments Nrf2-HO-1 pathways and anti-inflammatory properties in macrophages

Next, we investigated the effects of 4-oxoDHA on macrophages. The murine macrophage cell line (RAW 264.7) was treated with DHA, 4-HDHA, 4-oxoDHA (0.01–1 µM), or vehicle (N/C) for 6 h at 37 °C. The lysates were processed for western blot analysis to determine HO-1 expression. Treatment with 4-oxoDHA significantly enhanced HO-1 protein expression, reaching approximately 2.5-fold induction by 1 µM 4-oxoDHA (Figure 3a). Next, we investigated the activation of Nrf2, an upstream transcription factor for HO-1. Treatment with 4-oxoDHA significantly augmented the protein levels of Nrf2 (Figure 3b) and the DNA-binding activity of Nrf2 (Supplementary Figure 1).

**Figure 3.**
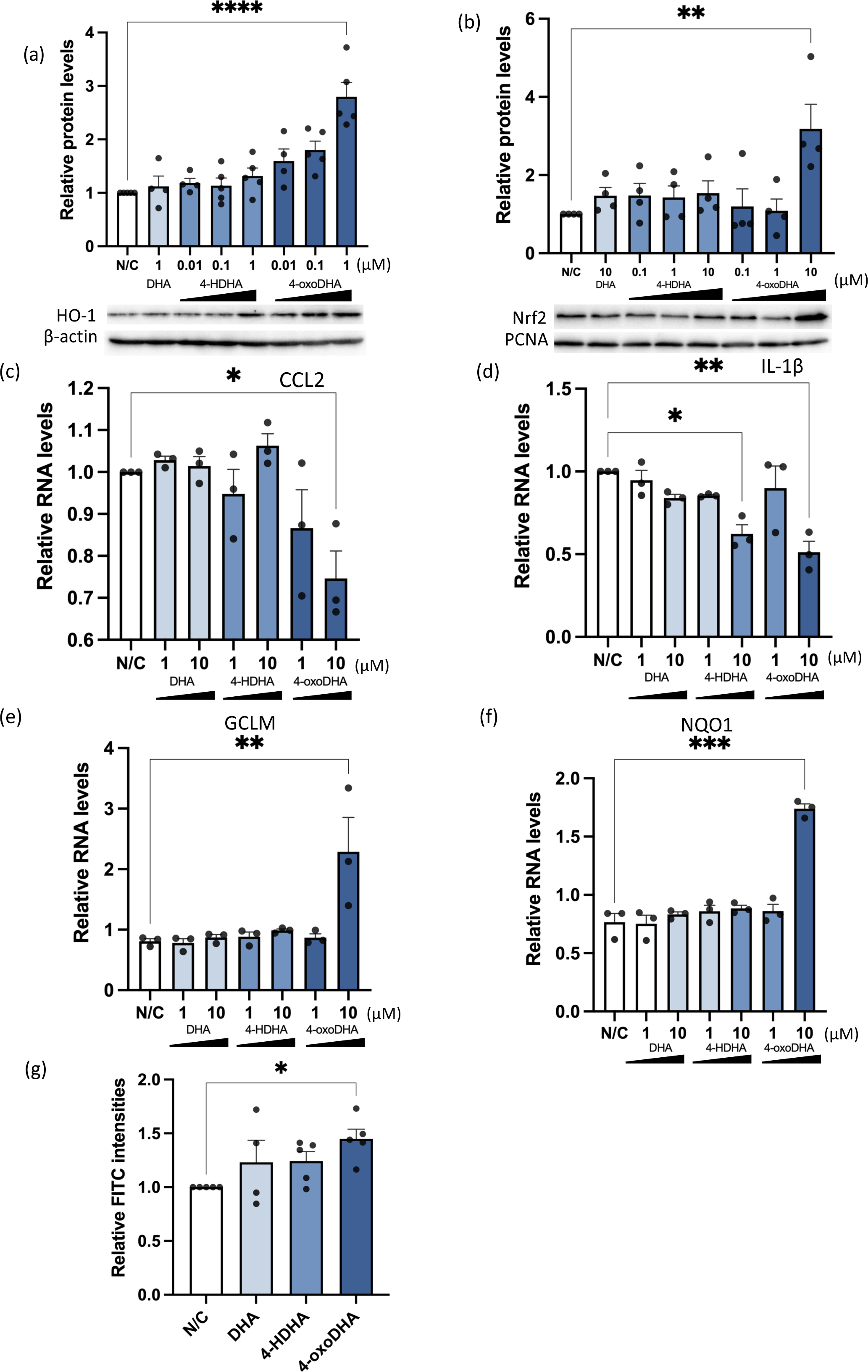
4-oxoDHA augments Nrf2-HO-1 pathways and anti-inflammatory properties in macrophages. (a) RAW 264.7 cells were treated with DHA, 4-HDHA, and 4-oxoDHA (0.01–1 µM) or vehicle (N/C) for 6h at 37°C. Lysates were processed for western blot analysis to determine HO-1 expression. β-actin was used as the internal standard. Data are expressed as fold change compared to vehicle, mean±s.e. (n=4–5). **** indicates p<0.0001. (b) RAW 264.7 cells were incubated with Kdo2 for 1h, followed by treatment of DHA (10 µM), 4-HDHA (0.1–10 µM), and 4-oxoDHA (0.1–10 µM) for 2h at 37°C. Nuclear fractions were processed for western blot analysis to determine Nrf2 expression. PCNA was used as the internal standard. Data are expressed as fold change compared to vehicle, mean±s.e. (n=4). ** indicates p<0.01. (c-f) RAW 264.7 cells were incubated with Kdo2 for 1h, followed by treatment with 1–10 µM of DHA, 4-HDHA, and 4-oxoDHA for 2h at 37°C. CCL2 (c), IL1-β (d), GCLM (e), and NQO1(f) mRNA were analyzed using real-time PCR. Data are expressed as fold change compared to vehicle, mean±s.e. (n=3). *, **, *** indicate p<0.05, p<0.01, p<0.001. (g) RAW 264.7 cells were incubated with 1 µM of DHA, 4-HDHA, and 4-oxoDHA for 6h at 37°C, and phagocytosis was evaluated with FITC-labeled zymosan particles. Data are expressed as fold change compared to vehicle, mean±s.e. (n=4). * indicates p<0.05.

Next, we investigated cytokine production in macrophages. RAW 264.7 cells were incubated with Kdo2 (0.5 μg/mL) for 1 h, followed by treatment with 1–10 µM of DHA, 4-HDHA, and 4-oxoDHA for 2 h at 37 °C. Treatment with 4-oxoDHA significantly suppressed inflammatory cytokines, including C-C motif chemokine (CCL) 2 (Figure 3c) and interleukin (IL)-1ß (Figure 3d), induced by Kdo2. Treatment with 4-oxoDHA also enhanced the mRNA expression of glutamate-cysteine ligase modifier subunit (GCLM) (Figure 3e) and NAD(P)H quinone dehydrogenase 1 (NQO1) (Figure 3f), which have anti-inflammatory properties.

Furthermore, we focused on the phagocytosis of macrophages, as phagocytosis activity is a critical function of macrophages for the resolution of inflammation (26). We treated RAW 264.7 cells with 1 µM DHA, 4-HDHA, and 4-oxoDHA for 6 h at 37 °C and evaluated phagocytosis with FITC-labeled zymosan particles. Treatment with 4-oxoDHA significantly enhanced phagocytosis (Figure 3g). These results indicate that 4-oxoDHA augments the Nrf2-HO-1 pathway and upregulates anti-inflammatory and pro-resolution functions in the macrophages.

### Noradrenaline regulates 5-LO protein expression via dopamine D2-like receptors in neutrophils

Given that 4-oxoDHA, regulating the anti-inflammatory functions of endothelial cells and macrophages, was reduced under stressed conditions, we hypothesized that specific blood cells produce 4-oxoDHA and that stress hormones regulate its production. First, heparin-treated whole blood (100 µL) was stimulated by specific stress hormones (adrenaline, noradrenaline, dopamine, and cortisol). Noradrenaline stimulation (10–1000 µM) for 1 h significantly suppressed the production of 4-HDHA and 4-oxoDHA in whole human blood (Figure 4a). Notably, postganglionic sympathetic fibers release noradrenaline under stressed conditions (12, 27).

**Figure 4.**
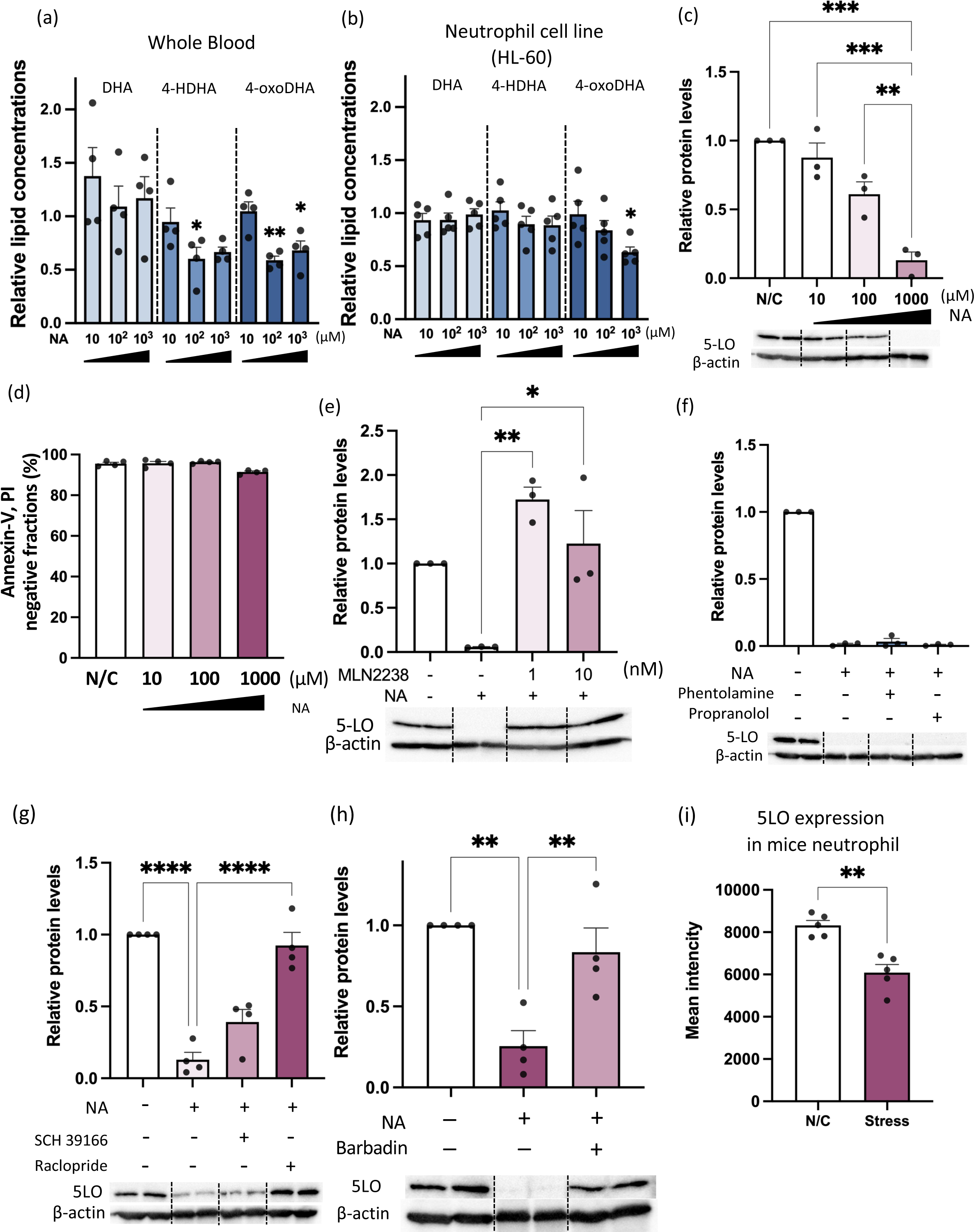
Noradrenaline regulates 5-LO protein expression via dopamine D2-like receptors in neutrophils. (a) Whole blood was stimulated with noradrenaline (NA) or vehicle for 1h at 37°C. Fatty acid metabolites were analyzed using LC/MS/MS. Data are expressed as fold change compared to the vehicle-treated group, mean±s.e. (n=3). *, ** indicate p<0.05, p<0.01 compared to vehicle. (b) Neutrophil cell line (HL-60) was treated with noradrenaline or vehicle for 2.5h at 37°C. Fatty acid metabolites were analyzed as above. * indicate p<0.05 compared to vehicle. (c) HL-60 cells were treated with noradrenaline for 2.5h at 37°C. Cell lysates were processed for western blot analysis to determine 5-LO expression. β-actin was used as the internal standard. Data are expressed as fold change compared to control, mean±s.e. (n=3). **, *** indicate p<0.01, p<0.001. (d) HL-60 cells were treated with noradrenaline for 2.5h at 37°C. Annexin-V and PI negative fractions are shown as percent of total cells, mean±s.e. (n=4) (e) HL-60 cells were treated with noradrenaline (1000 µM) for 2.5h at 37°C, with and without 1h pretreatment with MLN 2238 (proteasome inhibitor). Cell lysates were processed for western blot analysis to determine 5-LO expression. β-actin was used as the internal standard. Data are expressed as fold change compared to control, mean±s.e. (n=3). *, ** indicate p<0.05, p<0.01. (f) HL-60 cells were treated with noradrenaline (1000 µM) for 2.5h at 37°C, with and without 1-h pretreatment with phentolamine (alpha receptor inhibitor, 150 nM) and propranolol (beta receptor inhibitor, 120 nM). Cell lysates were processed for western blot analysis to determine 5-LO expression. β-actin was used as the internal standard. Data are expressed as fold change compared to control, mean±s.e. (n=4). (g) HL-60 cells were treated with noradrenaline (1000 µM) for 2.5h at 37°C, with or without pretreatment with SCH39166 (D1-like receptor antagonist, 10 nM) or raclopride (D2-like receptor antagonist, 20 nM). Data are expressed as fold change compared to control, mean±s.e. (n=4). **** indicate p<0.0001. (h) HL-60 cells were treated with noradrenaline (1000 µM) for 2.5h at 37°C, with and without 1-h pretreatment with Barbadian (β-arrestin inhibitor, 100 µM). Cell lysates were processed for western blot analysis to determine 5-LO expression. β-actin was used as the internal standard. Data are expressed as fold change compared to control, mean±s.e. (n=4). ** indicates p<0.01. (i) Mice were treated with or without stress by physical restriction (2h) for two consecutive days (as in Figure 1b). On day 3, 5-LO protein expression in neutrophils was monitored using flow cytometry. Data are expressed as mean fluorescent intensity, mean±s.e. (n=5). ** indicate p<0.01.

Next, we investigated which whole blood cells were responsible for the production of 4-HDHA and 4-oxoDHA. To this end, we employed neutrophil-like (HL60), monocyte-like (RAW 264.7), T lymphocyte-like (EL4), B lymphocyte-like (Ba/F3), and megakaryocyte cell lines (Meg-1) and profiled lipid metabolite production with or without noradrenaline treatment (data not shown). We found that the neutrophil-like cell line (HL60) biosynthesized 4-HDHA and 4-oxoDHA from DHA and that noradrenaline treatment reduced 4-oxoDHA levels by 30–40% (Figure 4b). The effect of noradrenaline treatment had the same tendency as found in human plasma under a depressive state (Table 1) and mouse plasma after 2 h of restraint stress (Figure 1a).

Because 4-oxoDHA is produced by 5-LO from DHA, we hypothesized that noradrenaline might regulate 5-LO expression in neutrophils. We investigated the mRNA and protein expression of 5-LO in HL60 cells after noradrenaline stimulation. HL60 cells were treated with noradrenaline (10–1000 μM) for 2.5 h at 37 °C, and cell lysates were processed for western blot analysis to determine 5-LO expression. The protein expression of 5-LO reduced by ∼40% with 100 µM noradrenaline and ∼90% with 1000 µM noradrenaline (Figure 4c), whereas the mRNA expression of 5-LO did not show any significant changes (Supplementary Figure 2). Stimulation with cortisol, also known as a stress hormone, did not significantly alter the expression of 5-LO proteins in HL60 cells (Supplementary Figure 3). We monitored annexin-V and PI double-negative fractions to exclude possible cellular toxicity induced by relatively high concentrations of noradrenaline. These fractions were not significantly altered by 10–1000 µM noradrenaline (Figure 4d).

We investigated how noradrenaline regulates 5-LO protein expression in neutrophils. Noradrenaline-derived suppression of 5-LO protein was rescued by MLN2238 (proteasome inhibitor; 1–10nM; 1 h before noradrenaline stimulation; Figure 4e), suggesting that the ubiquitin-proteasome system degraded the 5-LO protein. Next, we used selective agonists for adrenergic α receptors (Supplementary Figure 4a) and selective agonists for adrenergic β receptors (Supplementary Figure 4b). We found that none of these compounds suppressed 5-LO protein expression in neutrophils. Moreover, specific antagonists for each α- and β-adrenergic receptor failed to rescue the noradrenaline-derived suppression of the 5-LO protein expression (Figure 4f). We treated neutrophils with dopamine receptor antagonists (SCH39166 for D1-like receptors and raclopride for D2-like receptors), followed by noradrenaline stimulation, and found that raclopride completely recovered the noradrenaline-induced suppression of 5-LO protein expression in neutrophils (Figure 4g). We also employed selective dopamine receptor agonists (SKF81297 for D1-like receptors and quinpirole for D2-like receptors) and found no suppression of 5-LO protein expression in the neutrophils (Supplementary Figure 5).

Finally, Barbadian (β arrestin-2 inhibitor) treatment almost completely rescued noradrenaline-derived suppression of 5-LO (Figure 4h), indicating a relatively high concentration of noradrenaline-activated dopamine D2-like receptors via β arrestin-2 dependent signaling pathways and reduced 5-LO protein levels via the ubiquitin-proteasome system. We also measured 5-LO protein expression in neutrophils from restraint stress model mice. The mean intensity of 5-LO protein expression in neutrophils was ∼25% lower in the stressed group than in the control group (Figure 4i), indicating that 5-LO protein expression in neutrophils is also regulated in an in vivo restraint stress mouse model.

## Discussion

Our study reveals that stress conditions lead to a reduction in circulating levels of 4-oxoDHA, which is regulated by 5-LO in neutrophils. We found that stress triggers the degradation of 5-LO in neutrophils through the proteasome system in the dense distribution of noradrenaline within the bone marrow micro-compartment. Additionally, we discovered that dopamine D2-like receptor activation via the β-arrestin pathway is involved in the regulation of 5-LO. The decreased levels of circulating 4-oxoDHA resulted in the downregulation of the Nrf2-HO-1 anti-inflammatory axis and an increase in ICAM-1 expression, vascular permeability, and vascular remodeling (see Figure 5).

**Figure 5.**
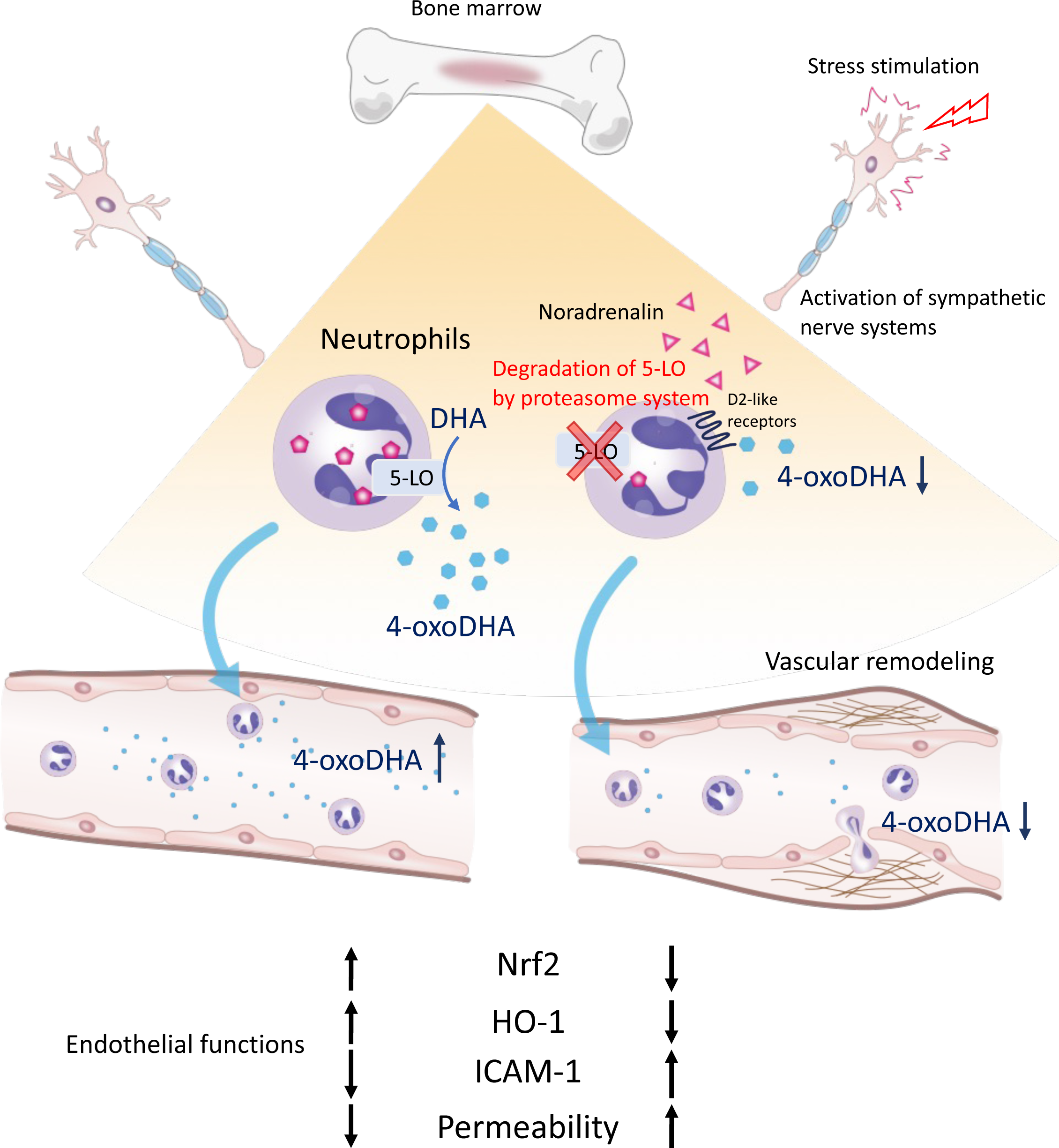
Circulating 4-oxoDHA is regulated by 5-LO expression in neutrophils and activates Nrf2 pathways, which maintain endothelial cell functions. However, under a stressed condition, a high concentration of noradrenaline is secreted in the bone marrow compartment, and D2-like receptor activation degrades 5-LO via the proteasome system in neutrophil, resulting in reduced levels of circulating 4-oxoDHA. This reduces Nrf2-HO-1 anti-inflammatory axis, resulting in increased ICAM-1 expression, vascular permeability, and remodeling.

Dietary consumption of omega-3 fatty acid-rich fish oil has been associated with various health benefits, including improved neurocognitive function (28), enhanced insulin resistance in individuals with diabetes (29), reduced cardiovascular event incidence (30, 31), and decreased inflammation (32). However, the precise molecular mechanisms underlying the beneficial effects of omega-3 fatty acids are not fully understood. Recent evidence suggests that the resolution of acute inflammation and the restoration of homeostasis involve active and highly regulated processes, termed programmed resolution, which are coordinated by bioactive lipid mediators derived from omega-3 fatty acids (33, 34). Nonetheless, little is known about the presence of functional lipid metabolites corresponding to stress-induced systemic inflammation in the bloodstream. Omega-3 PUFAs are metabolized and enzymatically converted into specialized pro-resolving mediators (SPMs), such as lipoxins, resolvins, protectins, and maresins (33). Enzymes like lipoxygenases (LOs), cytochrome p450, and cyclooxygenase (COX) play a role in the synthesis of bioactive anti-inflammatory lipids. Dehydrogenases convert the hydroxy group of these lipids into an oxo group, which imparts electrophilicity. Electrophilic fatty acids act through various signaling mechanisms, including noncovalent binding to receptors and covalent binding to target proteins and receptors (35, 36). Notably, 4-oxoDHA activates PPARγ(37) and regulates electrophile-sensitive signaling pathways, such as the Keap1/Nrf2-dependent antioxidant response. In the case of Nrf2/Keap1, electrophilic fatty acids induce the release and nuclear translocation of Nrf2 by modifying a specific cysteine residue on the Nrf2 inhibitor Keap1. Once in the nucleus, Nrf2 binds to antioxidant response elements (AREs) and activates the transcription of protective genes. Our study demonstrates that 4-oxoDHA increases Nrf2 protein expression (see Figure 2d, 3b) and enhances ARE-binding activities (see Supplementary Figure 1), subsequently inducing HO-1 (see Figure 2b, 2c, 3a) and glutathione biosynthetic enzymes (see Figure 3c).

Given that stress conditions are known to exacerbate vascular diseases, we utilized a mouse model of vascular remodeling that leads to stenotic lesions. Endothelial cells and macrophages have been shown to play critical roles in the pathogenesis of vascular diseases (38, 39). The induction of adhesion molecules by endothelial cells initiates the rolling, adhesion, and infiltration of leukocytes (40). Additionally, the loss of endothelial barrier function contributes to the extravasation of leukocytes (41). Macrophages regulate peripheral tissue homeostasis by phagocytosing apoptotic tissue debris and neutrophils (42). In our study, 4-oxoDHA reduced the expression of adhesion molecules in the mouse vascular remodeling model (see Figure 1g, 1h) and in an in vitro model (see Figure 2f, 2g). Furthermore, 4-oxoDHA enhanced macrophage phagocytosis (see Figure 3g) and exhibited anti-inflammatory properties (see Figure 3c, d, e, f).

While our study suggests that reduced levels of circulating 4-oxoDHA could serve as a novel biomarker for stress (see Table 1), we acknowledge that our study design was cross-sectional and, therefore, unable to establish a cause-and-effect relationship. Nevertheless, we consistently observed reduced circulating 4-oxoDHA levels in healthy volunteers experiencing depressive states and mice subjected to restraint stress. These findings suggest that specific stress-induced reactions may be conserved across species. Our study primarily focused on the sympathetic nervous system, as the bone marrow receives rich innervation from sympathetic nerves involved in regulating hematopoietic homeostasis (12, 27). Noradrenaline, secreted by sympathetic nerve terminals within the bone marrow, exhibits a dense distribution within the micro-compartment (21). Importantly, the plasma concentration of noradrenaline is approximately 1-2 nM (43), suggesting that noradrenaline can selectively regulate neutrophils within specific microenvironments like the bone marrow.

We concluded that high levels of noradrenaline, acting through specific receptors other than adrenergic receptors, lead to the reduction of 5-LO protein expression in neutrophils. This conclusion is supported by our observations that selective agonists for α and β adrenergic receptors failed to reduce 5-LO protein expression (see Supplementary Figure 4) and that selective antagonists for α and β receptors did not rescue the noradrenaline-induced reduction in 5-LO protein (see Figure 4f). Notably, noradrenaline has been shown to act as a potent agonist for all D2-like receptor subtypes (44). Furthermore, we found that the D2-like receptor antagonist raclopride effectively rescued the noradrenaline-induced reduction in 5-LO protein (see Figure 4g), and the β-arrestin inhibitor Barbadian maintained 5-LO protein expression even under noradrenaline stimulation (see Figure 4h). These results demonstrate that noradrenaline regulates 5-LO protein expression in neutrophils through dopamine D2-like receptors via the β-arrestin pathway.

Collectively, our findings provide evidence for a novel stress-induced vascular inflammation pathway. Reduced circulating levels of 4-oxoDHA lead to compromised Nrf2-ARE-related anti-inflammatory functions in endothelial cells and macrophages, resulting in vascular inflammation. These neuro-lipid metabolite interactions under stress conditions may serve as biomarkers for stress and offer a novel target for preventing systemic neuroinflammation.

## Acknowledgments

We are grateful to Dr. Hirotaka Nagai and Dr. Ryota Shinohara for their helpful discussions. This study was partly supported by grants from AMED (JP22gm0910012 and JP22wm0425001 to T.F.) and KAKENHI (18H05429 and 21H04812 to T.F. and 20K08422 to M.S.).

## Supplementary Material (Tables and Figures)

**Supplementary Figure 1**

RAW 264.7 cells were incubated with Kdo2 (0.5 μg/mL) for 1h, followed by treatment with 10 µM DHA, 4-HDHA, and 4-oxoDHA for 2h at 37°C. The DNA binding activity of Nrf2 was significantly augmented by 4-oxoDHA treatment. Data are expressed as fold induction of DNA-binding activity compared to vehicle (N/C), mean±s.e. (n=5). * indicates p<0.05.

**Supplementary Figure 2**

HL-60 cells were treated with noradrenaline (10–1000 μM) for 2.5h at 37°C, and 5-LO mRNA expression levels were analyzed using real-time PCR. The 5-LO mRNA expression did not show any significant changes. Data are expressed as fold-change compared to vehicle (N/C), mean±s.e. (n=5).

**Supplementary Figure 3**

HL-60 cells were treated with cortisol (1–10 μM) for 2.5h at 37°C, and 5-LO protein expression levels were analyzed using western blotting. The 5-LO protein expression did not show any significant changes. Data are expressed as fold-change compared to vehicle (N/C), mean±s.e. (n=3).

**Supplementary Figure 4**

HL-60 cells were treated with noradrenaline (1000 μM) and (a) selective adrenergic α receptor agonists (cirazoline and azepexole, 100 µM each) or (b) selective adrenergic β receptor agonists (dobutamine, clenbuterol, and BRL37344, 100 µM each) for 2.5h at 37°C, and 5-LO protein expression levels were analyzed using western blotting. Neither selective adrenergic α-receptor agonists (a) nor selective adrenergic β-receptor agonists (b) reduced 5-LO protein expression, whereas noradrenaline treatment significantly reduced 5-LO protein expression. Data are expressed as fold-change compared to vehicle (N/C) and mean±s.e. (n=4). * indicates p<0.05.

**Supplementary Figure 5**

HL-60 cells were treated with SKF81297 (D1-like receptor agonist, 10 µM) or quinpirole (D2-like receptor agonist, 10 µM) for 2.5h at 37°C, and 5-LO protein expression levels were analyzed using western blotting. Neither the D1-like receptor agonist nor the D2-like receptor agonist reduced 5-LO protein expression, whereas noradrenaline (1000 μM) or dopamine (DOA, 500 µM) treatment significantly reduced 5-LO protein expression. Data are expressed as fold-change compared to vehicle (N/C) and mean±s.e. (n=4). **** indicates p<0.0001.

## Highlights

- Our study reveals that stress-induced reduction in circulating levels of a specific omega-3 fatty acid metabolite, 4-oxoDHA, contributes to vascular inflammation.
- We have identified a novel pathway that explains how stress promotes systemic vascular inflammation by regulating omega-3 fatty acid metabolites in the circulation.
- Our findings provide new evidence for the role of 4-oxoDHA in maintaining Nrf2-ARE-related anti-inflammatory functions in endothelial cells and macrophages.

## Notes

### Competing Interest Statement

The authors have declared no competing interest.

## References

1. Yaribeygi H, Panahi Y, Sahraei H, Johnston TP, Sahebkar A. The impact of stress on body function: A review. EXCLI J. 2017;16:1057–72.

2. McEwen BS, Bowles NP, Gray JD, Hill MN, Hunter RG, Karatsoreos IN, et al. Mechanisms of stress in the brain. Nat Neurosci. 2015;18(10):1353–63.

3. Black PH. The inflammatory response is an integral part of the stress response: Implications for atherosclerosis, insulin resistance, type II diabetes and metabolic syndrome X. Brain Behav Immun. 2003;17(5):350–64.

4. Yusuf S, Hawken S, Ounpuu S, Dans T, Avezum A, Lanas F, et al. Effect of potentially modifiable risk factors associated with myocardial infarction in 52 countries (the INTERHEART study): case-control study. Lancet. 2004;364(9438):937–52.

5. Rosengren A, Hawken S, Ounpuu S, Sliwa K, Zubaid M, Almahmeed WA, et al. Association of psychosocial risk factors with risk of acute myocardial infarction in 11119 cases and 13648 controls from 52 countries (the INTERHEART study): case-control study. Lancet. 2004;364(9438):953–62.

6. Chumaeva N, Hintsanen M, Ravaja N, Puttonen S, Heponiemi T, Pulkki-Raback L, et al. Interactive effect of long-term mental stress and cardiac stress reactivity on carotid intima-media thickness: the Cardiovascular Risk in Young Finns study. Stress. 2009;12(4):283–93.

7. Lagraauw HM, Kuiper J, Bot I. Acute and chronic psychological stress as risk factors for cardiovascular disease: Insights gained from epidemiological, clinical and experimental studies. Brain Behav Immun. 2015;50:18–30.

8. Flower RJ. Prostaglandins, bioassay and inflammation. Br J Pharmacol. 2006;147 Suppl 1(Suppl 1):S182–92.

9. Samuelsson B. Role of basic science in the development of new medicines: examples from the eicosanoid field. J Biol Chem. 2012;287(13):10070–80.

10. Dinarello CA, Simon A, van der Meer JW. Treating inflammation by blocking interleukin-1 in a broad spectrum of diseases. Nat Rev Drug Discov. 2012;11(8):633–52.

11. Serhan CN, Savill J. Resolution of inflammation: the beginning programs the end. Nat Immunol. 2005;6(12):1191–7.

12. Mendez-Ferrer S, Lucas D, Battista M, Frenette PS. Haematopoietic stem cell release is regulated by circadian oscillations. Nature. 2008;452(7186):442-7.

13. Gross RW, Jenkins CM, Yang J, Mancuso DJ, Han X. Functional lipidomics: the roles of specialized lipids and lipid-protein interactions in modulating neuronal function. Prostaglandins Other Lipid Mediat. 2005;77(1-4):52–64.

14. Tsui-Pierchala BA, Encinas M, Milbrandt J, Johnson EM, Jr. Lipid rafts in neuronal signaling and function. Trends Neurosci. 2002;25(8):412–7.

15. Parekh A, Smeeth D, Milner Y, Thure S. The Role of Lipid Biomarkers in Major Depression. Healthcare (Basel). 2017;5(1).

16. Liao Y, Xie B, Zhang H, He Q, Guo L, Subramanieapillai M, et al. Efficacy of omega-3 PUFAs in depression: A meta-analysis. Transl Psychiatry. 2019;9(1):190.

17. Zung WW. A Self-Rating Depression Scale. Arch Gen Psychiatry. 1965;12:63–70.

18. Thaker PH, Han LY, Kamat AA, Arevalo JM, Takahashi R, Lu C, et al. Chronic stress promotes tumor growth and angiogenesis in a mouse model of ovarian carcinoma. Nat Med. 2006;12(8):939–44.

19. Cui B, Luo Y, Tian P, Peng F, Lu J, Yang Y, et al. Stress-induced epinephrine enhances lactate dehydrogenase A and promotes breast cancer stem-like cells. J Clin Invest. 2019;129(3):1030–46.

20. Kumar A, Lindner V. Remodeling with neointima formation in the mouse carotid artery after cessation of blood flow. Arterioscler Thromb Vasc Biol. 1997;17(10):2238–44.

21. Kawano Y, Fukui C, Shinohara M, Wakahashi K, Ishii S, Suzuki T, et al. G-CSF-induced sympathetic tone provokes fever and primes antimobilizing functions of neutrophils via PGE2. Blood. 2017;129(5):587–97.

22. Colas RA, Shinohara M, Dalli J, Chiang N, Serhan CN. Identification and signature profiles for pro-resolving and inflammatory lipid mediators in human tissue. Am J Physiol Cell Physiol. 2014;307(1):C39–54.

23. Bolger AM, Lohse M, Usadel B. Trimmomatic: a flexible trimmer for Illumina sequence data. Bioinformatics. 2014;30(15):2114–20.

24. Patro R, Duggal G, Love MI, Irizarry RA, Kingsford C. Salmon provides fast and bias-aware quantification of transcript expression. Nat Methods. 2017;14(4):417-+.

25. Ge SX, Son EW, Yao RN. iDEP: an integrated web application for differential expression and pathway analysis of RNA-Seq data. Bmc Bioinformatics. 2018;19.

26. Kourtzelis I, Hajishengallis G, Chavakis T. Phagocytosis of Apoptotic Cells in Resolution of Inflammation. Front Immunol. 2020;11:553.

27. Katayama Y, Battista M, Kao WM, Hidalgo A, Peired AJ, Thomas SA, et al. Signals from the sympathetic nervous system regulate hematopoietic stem cell egress from bone marrow. Cell. 2006;124(2):407–21.

28. Morris MC, Evans DA, Tangney CC, Bienias JL, Wilson RS. Fish consumption and cognitive decline with age in a large community study. Arch Neurol-Chicago. 2005;62(12):1849–53.

29. Fedor D, Kelley DS. Prevention of insulin resistance by n-3 polyunsaturated fatty acids. Curr Opin Clin Nutr Metab Care. 2009;12(2):138–46.

30. Yokoyama M, Origasa H, Matsuzaki M, Matsuzawa Y, Saito Y, Ishikawa Y, et al. Effects of eicosapentaenoic acid on major coronary events in hypercholesterolaemic patients (JELIS): a randomised open-label, blinded endpoint analysis. Lancet. 2007;369(9567):1090–8.

31. Bhatt DL, Steg PG, Miller M, Brinton EA, Jacobson TA, Ketchum SB, et al. Cardiovascular Risk Reduction with Icosapent Ethyl for Hypertriglyceridemia. N Engl J Med. 2019;380(1):11–22.

32. Duda MK, O’Shea KM, Tintinu A, Xu W, Khairallah RJ, Barrows BR, et al. Fish oil, but not flaxseed oil, decreases inflammation and prevents pressure overload-induced cardiac dysfunction. Cardiovasc Res. 2009;81(2):319–27.

33. Serhan CN. Pro-resolving lipid mediators are leads for resolution physiology. Nature. 2014;510(7503):92–101.

34. Tabas I, Glass CK. Anti-inflammatory therapy in chronic disease: challenges and opportunities. Science. 2013;339(6116):166–72.

35. Rudolph TK, Freeman BA. Transduction of redox signaling by electrophile-protein reactions. Sci Signal. 2009;2(90):re7.

36. Groeger AL, Cipollina C, Cole MP, Woodcock SR, Bonacci G, Rudolph TK, et al. Cyclooxygenase-2 generates anti-inflammatory mediators from omega-3 fatty acids. Nat Chem Biol. 2010;6(6):433–41.

37. Itoh T, Fairall L, Amin K, Inaba Y, Szanto A, Balint BL, et al. Structural basis for the activation of PPARgamma by oxidized fatty acids. Nat Struct Mol Biol. 2008;15(9):924–31.

38. Rozanski A, Blumenthal JA, Kaplan J. Impact of psychological factors on the pathogenesis of cardiovascular disease and implications for therapy. Circulation. 1999;99(16):2192–217.

39. Kivimaki M, Steptoe A. Effects of stress on the development and progression of cardiovascular disease. Nat Rev Cardiol. 2018;15(4):215–29.

40. Ley K, Laudanna C, Cybulsky MI, Nourshargh S. Getting to the site of inflammation: the leukocyte adhesion cascade updated. Nat Rev Immunol. 2007;7(9):678–89.

41. Sluiter TJ, van Buul JD, Huveneers S, Quax PHA, de Vries MR. Endothelial Barrier Function and Leukocyte Transmigration in Atherosclerosis. Biomedicines. 2021;9(4).

42. Fredman G, Hellmann J, Proto JD, Kuriakose G, Colas RA, Dorweiler B, et al. An imbalance between specialized pro-resolving lipid mediators and pro-inflammatory leukotrienes promotes instability of atherosclerotic plaques. Nat Commun. 2016;7:12859.

43. Levin BE, Natelson BH. The relation of plasma norepinephrine and epinephrine levels over time in humans. J Auton Nerv Syst. 1980;2(4):315–25.

44. Sanchez-Soto M, Bonifazi A, Cai NS, Ellenberger MP, Newman AH, Ferre S, et al. Evidence for Noncanonical Neurotransmitter Activation: Norepinephrine as a Dopamine D2-Like Receptor Agonist. Mol Pharmacol. 2016;89(4):457–66.

